# shRNA mediated inhibition of Cdc42 gene expression in Calu-6, lung cancer cells

**DOI:** 10.1101/161612

**Authors:** Zohreh Ghambari, Mohammad Nabiuni, Hanieh Jalali, Latifeh Karimzadeh

**Affiliations:** Department of Cell and Molecular Biology, Faculty of Biological Sciences, Kharazmi University, Tehran, Iran; Department of Animal Biology, Faculty of Biological Science, Kharazmi University, Tehran, Iran

**Keywords:** Lentivirus, Lung cancer, Gene therapy, Cdc42, shRNA

## Abstract

**Background Information:** RNAi technique as a new strategy in gene therapy is the effective gene silencing method. Cdc42 is a member of Rho GTPases involving in lung cancer cells migration and proliferation. In the present study, targeting and inhibiting the Cdc42 expression in Calu-6 cells was investigated. Recombinant lentiviral particles were produced by co-transfection of pMD2.G, psPAX2 and pGFP-C-shLenti plasmid in 293T packaging cells. Calu-6 cells were transduced by recombinant lentiviruses using polybrene. GFP-fluorescence microscopy and MTT assay were used to assess the Calu-6 target cell transduction and rate of lentiviral transduced cells proliferation, respectively. Real time PCR was performed to compare the expression of Cdc42 gene before and after shRNA delivery.

**Results:** GFP-fluorescence microscopy analysis showed that Calu-6 cells were successfully transduced with recombinant lentiviral expressing shRNA-Cdc42. The viability of transduced cells was reduced within 72, 96 and 120 hours of transduction process. Real time PCR analyze showed the significant reduction of Cdc42 gene expression. **Conclusions:** Lentiviral vectors may be reasonable tools for this gene delivery due to stable expression of silencing RNA. Inhibition of Cdc42 expression by lentiviral mediated shRNA delivery could be an effective method to inhibit proliferation of lung cancer cells. **Significance:** Over expression of Cdc42 gene have been seen in lung cancer. This make Cdc42 gene a key target in treatment of cancer. On the other hand, gene therapy is a proper method to modulate gene expression. It seems modulation of cdc42 gene expression by gene therapy accompany with proper vehicles creating hopes in treatment of lung cancer.

## 1 Introduction

Lung cancer is the most leading cause of cancer death in the worldwide (Dellaire et al. 2014). Surgery, chemotherapy and radiotherapy are three common lung cancer treatments and sometimes have been used in combination; these methods usually work in early stages of cancer and is not very efficient in late stages or metastatic form of lung cancer and have many adverse effects (Sakiragaoglu et al. 2013).

Gene therapy is a method for modifying the defective genome function. The earliest strategy of Gene therapy was providing a functional version of defective gene (Bolhassani & Saleh 2013). More recently, new strategies have been developed in gene therapy which manage cell pathways by target gene correction or disruption.(Humbert et al. 2012).

Using a short hairpin RNA (shRNA) as a part of RNAi technology can be used in cancer targeted therapy. In shRNA-based RNAi technology, sequence-specific gene shRNA as known Pri-shRNA is synthesized and processed by Drosha/DGCR8 complex in nucleus and transported into cytoplasm by Exportin 5 protein. In cytoplasm, pri-shRNA processed by dicer and incorporated into the RNA-interfering silencing complex (RISC) and activated specifically based on gene sequence, leading to targeted mRNA cleavage and degradation (Donald D. Rao et al. 2009).

Although gene therapy has been developed remarkably in last two decades, choosing a suitable vector for transfer of interest gene is still a challenge in gene therapy and efforts keep going on design a safer and long-term gene expression vectors. Among viral and non-viral vectors, Lentivirus mediated vectors have their own advantages for gene delivery purposes. Lentiviral vectors are a type of retrovirus that can infect both dividing and non-dividing cells. Lacking of genes required for self-replication in lentiviral vectors lead to their replicate defection and make them as safe vehicle for gene therapy trials. On the other hand, lentiviruses capability of stable transduction make them highly effective and suitable in gene therapy techniques (Song & Yang 2010).

Cdc42 is a member of Rho GTPases family whit a wide range function in the cell including filo podia formation, cell movement, and rearrangement of cytoskeleton, cell migration and proliferation. Upregulation of Cdc42 have been shown in some cancers such as non-small cell lung cancer, colorectal adenocarcinoma, melanoma, breast cancer and testis cancer. The function of Cdc42 gene as a key target in cancer therapy Elevated via collapse degradation of EGFR by c-Cbl and cause EGFR more function and increase cell proliferation (Stengel & Zheng 2011; Gómez Del Pulgar et al. 2008; Chou et al. 2003; Qadir et al. 2015).

In this investigation, we transferred Cdc42 sequence specified shRNA into lung cancer cell line Calu6 by lentiviral vector for stably production of inhibitory shRNA and the expression of Cdc42 gene as a suitable method and target in lung cancer therapy and Calu-6-transduced cells proliferation rate were assayed.

## 2 Materials and Methods

### 2.1 Cell lines and Culture

Calu-6 lung cancer cell line and 293T embryonic cell line were purchased from National Cell Bank of Iran (NCBI, Pasture Institute of Iran, Tehran). Cells were cultured in RPMI-1640 supplemented with 10% fetal bovine serum, 1% penicillin-streptomycin in a humidified atmosphere of 95% air and 5% CO2 at 37 °C. The medium was changed every 3 days.

### 2.2 Viral packaging and production

Transferring plasmid (tGFP-C-shLenti), possessing shRNA sequence against human Cdc42 gene and Green Fluorescent Protein (GFP) under the control of cytomegalovirus (CMV) promotor was purchased (Origene, China) (Figure 1).

**Figure 1:**
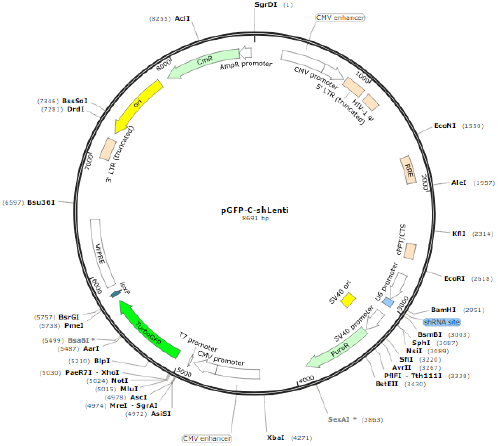
HuSH-29 lentiviral vector with chloramphenicol and puromycin resistance markers and a TurboGFP gene, for expressing an shRNA from the U6 promoter (http://www.snapgene.com).

Both shRNA Cdc42 sequence containing and non-containing recombinant lentiviruses were produced by transient transfection of 293T cells (3 × 10^6^ cell/ml) plated in 60-mm dishes. The tGFP-C-shLenti (10 μg), pMD2G, (10 μg, Addgene), and psPAX (10 μg, Addgene) plasmids were co-transfected into 293T cells using lipofectamine 2000 (Invitrogen, USA), according to the manufacturer’s instructions. Transfection was confirmed using GFP-positive cells observation under florescent microscopy (Micros, Austria). Images captured by USB DCam and iViewCap application. The culture supernatants containing virus particles were harvested by 48 and 72 hours post-transfection and filtered through 0.45-μm pore-size filters.

### 2.3 Lentiviral delivery to Calu-6 cells

To obtain the most efficient concentration of virus for transduction of Calu-6 cells, different dilutions of virus stock were added to cell culture medium and the percentage of GFP positive cells were determined by Fluorescence-activated cell sorting (FACS). For this purpose, 1x10^5^ calu-6 cells were seeded in 6 well plates and transduced by 0, 50, 125, 250 and 350 μl of lentiviral stock using Polybrene (Sigma Aldrich, UK) as transduction reagent. Forty eighth hours after transduction, transduced cells harvested and flow cytometry analysis was done to determine the rate of GFP expression.

### 2.4 Assessment of cytotoxicity

Cytotoxic effect of lentivirus infection on calu-6 cells was determined by 3-(4,5-dimethylthiazol-2-yl)-2,5-diphenyltetrazolium bromide (MTT) (Sigma Aldrich, UK) reduction assay in both shRNA containing and non-containing groups Which relays on the mitochondrial dehydrogenase activities. Briefly, lentiviruses in 1, 10, 50, 100 and 200 MOI was added to 6000 Calu-6 cells in 96-well plate. After 48 h, 10 ul of MTT (5mg/ml) solution was added to culture medium and incubation was done for 4 h. The dark blue formazan crystals were dissolved in 100 μl DMSO and absorbance was recorded at 490 nm. Percentage of viable cells was calculated as: OD test / OD control×100 formulas.

### 2.5 Antibiotic selection of infected cells

The log growth phase Calu-6 cells were seeded in 6-well plates in 1 × 10^5^ density and cultured overnight. For infection, the shRNA containing and non-containing lentiviruses were diluted in complete medium containing Polybrene (8 μg/mL) in 50 MOI and treated for 24 h at 37 °C. By the time of incubation, the virus containing medium was replaced with fresh medium and puromycin selection of infected cells was done; Minimal lethal dose of puromycin for Calu-6 cell line was determined during 2 days; For this aim, Calu-6 cell line treated by 0 (as control), 0.5-10 μg/ml of puromycin. Cells observed each day by invert microscope (Micros, Austria). GFP expression in survived cells was verified by fluorescent microscopy. Non-infected cells treated with puromycin used as control group for antibiotic selection. Images captured by USB DCam and iViewCap application.

### 2.6 Real-time PCR

To investigate the efficiency of Cdc42 mRNA knockdown by LV-shRNA transferring, real time PCR assay was performed. Briefly, total RNA was extracted from cells using RNA extraction kit (Thermo Scientific, USA) and converted to cDNA using cDNA Synthesis kit (Thermo Scientific, USA). Aliquots of cDNA were subjected to quantitative real-time PCR using Real-time PCR kit (TAKARA, USA). The mRNA levels were normalized to that of glyceraldehyde-3-phosphate dehydrogenase (GAPDH) housekeeping gene. The specific primer pairs were as follows: Cdc42, sense: 5′-GCCCGTGACCTGAAGGCTGTCA-3′; antisense: 5′-TGCTTTTAGTATGATGCCGACACCA-3′; GAPDH, sense: 5′-GGCATTGCTCTCAATGACAA -3′ reverse: 5′-TGTGAGGGAGATGCTCAGTG-3′. None transduce cells were subjected as control and LV-shRNA without Cdc42 silencing sequence infected cells were selected as negative control groups.

### 2.7 Proliferation assay

The effect of Cdc42 silencing on cell proliferation process was measured via MTT colorimetric assay. Lentivirus infected cells were seeded at a density of 2000 cells/well in 24-well plates. Experiments were designed as CON, NC and Lv-shRNACdc42F. After 24, 48, 72, 96 and 120 h incubation at 37 °C, 50 μL of sterile MTT dye (5 mg/mL; Sigma-Aldrich Corp) was added to each well and incubated for another 4 h at 37 °C. Medium was removed from the wells and dimethyl sulfoxide (DMSO) was added to each well for dissolving the formazan product. Spectrometric absorbance was measured at the wavelength of 490 nm.

### 2.8 Statistical analysis

All experiments were performed in triplicate. The values obtained from this study were expressed as mean ± standard deviation (SD). Statistically significant differences between groups were determined by One-way ANOVA using SPSS 16 software. Differences were considered statistically significant at P <0.05.

## 3 Results

### 3.1 Virus production in 293T cells

Recombinant lentivirus packaging and production was done in 293T cells and fluorescence microscopy observation showed that 24 h after infection, 80% of 293T Cells received virus production constructs and were expressing GFP marker (Figure 2).

**Figure 2.**
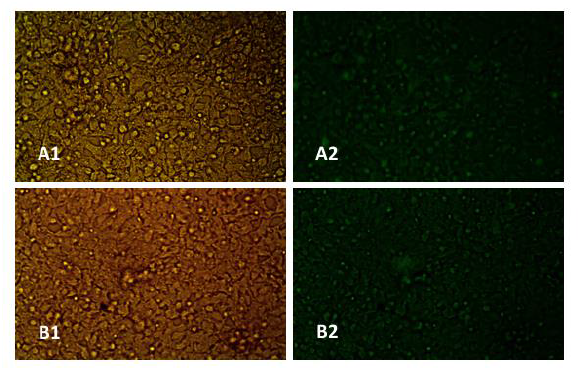
293T cells transfection. To produce recombinant lentivirus encoding shRNA, 293T cells co-transfected by psPAX2, pMD2G and transfer vector (tGFP-C-shLenti plasmid for lentivirus and control plasmid for control lentivirus) by lipofectamin 2000. 24 h after transfection, transfection efficiency observed by florescent microscopy. About 80% of cells transfected successfully. (A1) invert microscopy (A2) florescent microscopy of cells transfected by control plasmid. (B1) invert microscopy (B2) florescent microscopy of cells transfected by tGFP-C-shLenti plasmid. X400.

### 3.2 Viral transduction of Calu-6 cells

Treatment of Calu-6 cells with different dilutions of harvested virus showed that GFP expression in cells transduced by tGFP-C-shLenti encoding lentivirus was 12.7%, 15.30%, 11.35% and 2.10% in the presence of 350, 250, 125 and 50 μl of lentivirus stock solution respectively. For cells transduced by non shRNA expressing lentivirus, GFP expression percentage was 29.33%, 19.35, 8.75% and 3.34% respectively for 350, 250, 125 and 50 μl of virus stock (Figure 3). Based on these results, different MOIs calculated. Toxicity of lentiviruses (not shRNA) was assessed with MTT assay which showed no toxic effects on Calu-6 cells (Figure 4)

**Figure 3.**
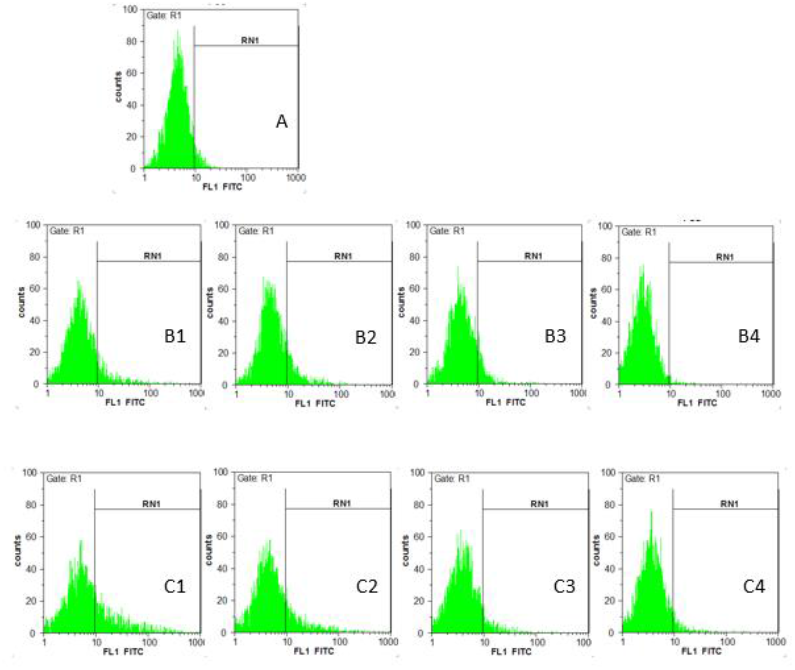
Determination of lentivirus stock titration. To determination of GFP gene expression percent in transduced cells by control and tGFP-C-shLenti encoding lentiviruses, Calu-6 cells treated by 0 (as a control), 50, 125, 250 and 350 μl of each lentivirus stock groups. GFP gene expression in cells transduced by tGFP-C-shLenti encoding lentivirus was (B1) 12.73%, %, (B2) 15.30, (B3) 11.35% and (B4) 2.10% and for cells transduced by control lentivirus was (C1) 29.33%, (C2) 19.35, (C3) 8.75% and (C4) 3.34% for 350, 250, 125 and 50 μl of lentivirus stock solution respectively.

**Figure 4.**
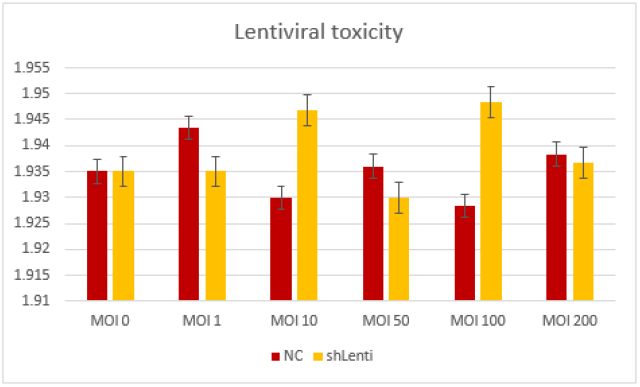
Assessment of recombinant lentivirus toxicity. To determine toxicity of generated lentiviruses (not shRNA) and their effect on assay results, Calu-6 cell line transduced by different MOI 0, 10, 50, 100 and 200 of Cdc42-shRNA encoding lentiviruse and control lentiviruse. MTT assay used to determine cytotoxicity of lentivirus stocks. Data shows no significant change in cell grows in different MOI of lentivirus groups.

### 3.3 Antibiotic selection of transduced cells

Calu-6 cells transduced by Cdc42-shRNA encoding lentivirus and control lentivirus purified by puromycin antibiotic. Results showed that after two days of antibiotic treatment, all cells died at 0.5 μg/ml of puromycin, then this dose was selected as minimal lethal dose of puromicyn against Calu6 cells (Figure 5). To remove all non-infected cells, antibiotic treatment continued for 4 weeks at 0.5 μg/ml μg/ml of puromycin which led to acquiring pure infected Calu6 cells (Figure 6).

**Figure 5.**
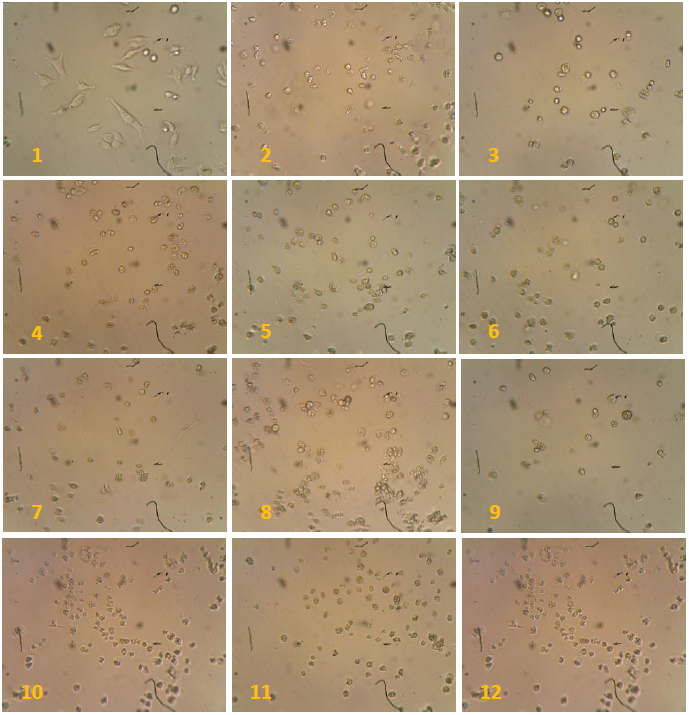
Treatment of Calu6 cells with puromycin to determine minimal lethal dose at 0.5-10 μg/ml. Observation under inverted microscopy at x250 Magnification. 1: control, 2-12: cell death under treatment with puromycin.

**Figure 6.**
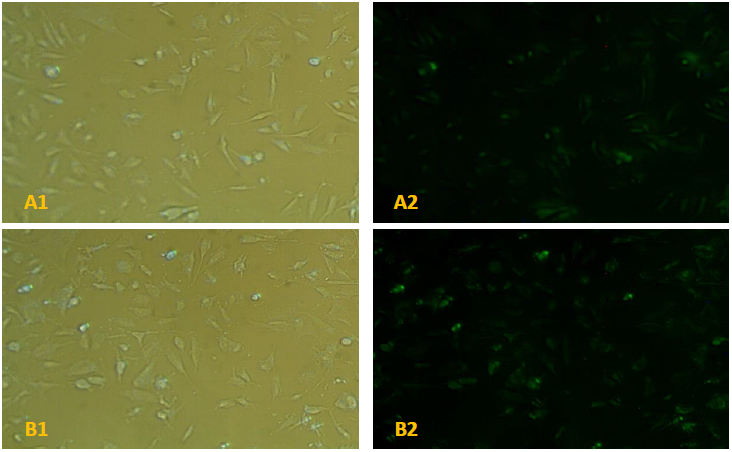
Purification of Calu-6 cellstransduced by lentiviruses 4 weeks after antibiotic treatment, only infected cells were alive and transduced cells were purified. (A1) invert microscopy (A2) florescent microscopy of cells infected by NC lentivirus. (B1) invert microscopy (B2) florescent microscopy of cells infected by tGFP-C-sh lentivirus. X400.

### 3.4 Cdc42 gene expression reduced in Calu-6 cells infected by tGFP-C-shLenti

To investigate the effect of shRNACdc42 on Cdc42 gene expression in calu-6 cells, real-time PCR assay performed and the expression of Cdc42 mRNA was compared between shRNACdc42 containing and non-containing cells. A Delta CT analysis indicated that the expression of Cdc42 gene in transduced cells was significantly decreased in compare to both control and negative control groups. Statistical analysis showed that the amount of Cdc42 mRNA was 0.25-fold in shRNACdc42 expressing cells compared to non-expressing cells (Figure 7).

**Figure 7.**
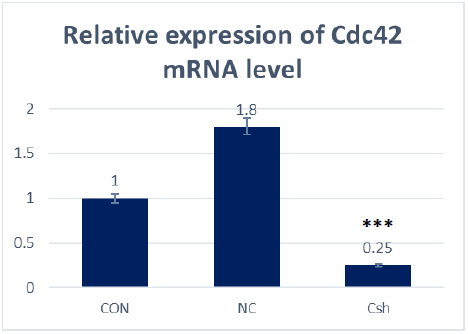
Expressions of Cdc42 mRNA in Calu-6 cells infected with Cdc42-shRNA encoding lentivirus. The expression levels of Cdc42 mRNA was assessed by qRT-PCR (2^-Δ Δ Ct^ method). 2^-Δ Δ Ct^ indicated reduction in relative expression level of Lv-shRNACdc42 group, compared with the CON group (***P<0.001) but indicated no Significant change in relative expression level of NC group compared with the CON group.

**Figure 8.**
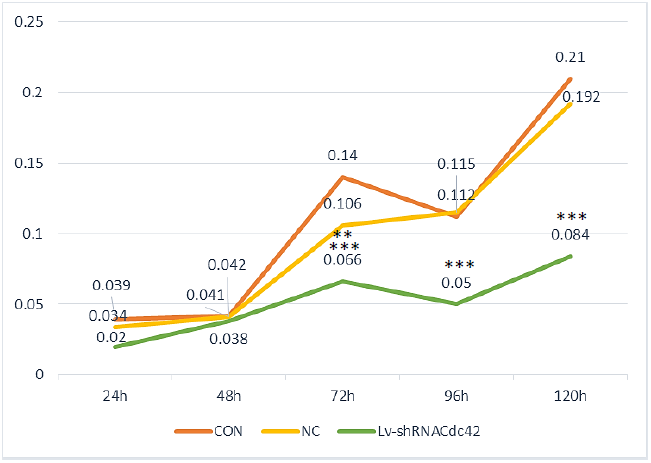
Assessment of calu-6 cell line proliferation. To investigate proliferation of Calu-6 cells transduced by Cdc42-shRNA encoding virus (Lv-shRNACdc42 cells), NC Calu-6 cells and CONT cells (non-transduced), cells seeded in 24 well plate in 2000 density. Proliferation rate of cells determined by MTT in 5 days. The diagram shows linear grows of cell lines. In 24 h and 48 h, significant differences don’t observed between grows rate of Lv-shRNACdc42 cells compare to NC cells and CON cells. In 72 h, significant differences observed in grows rate of Lv-shRNACdc42 cells compare to NC cells indicated by ** and P<0.01 and compare to CON cells by *** and P<0.001. In 96h and 120h, grows rate of Lv-shRNACdc42 cells compare to NC cells and CON cells was significant indicated by *** and P<0.001. Significant differences don’t observed between CON cells and NC cells.

### 3.5 Knockdown of Cdc42 inhibits proliferation of calu-6 cells

To determine whether down-regulation of Cdc42 affects cell growth in vitro, the capability of cell proliferation was assessed by MTT assay after Cdc42-shRNA encoding lentivirus infection. Compared with the CONT group and the NC group, cell proliferation in Cdc42-shRNA group is dramatically reduced suggesting that the knockdown of Cdc42 gene results in suppression of cell growth.

At the 24 and 48 h after seed cells, there was no significant reduction in grows rate of Lv-shRNACdc42 cells compare to CON and NC cells. Neither control shRNA transduced nor untransduced cells, demonstrated any change in viability over the two day time course. In day 3, reduce in Lv-shRNACdc42 cells grows was significant so that grows rate of Lv-shRNACdc42 cells compare to NC cells with two stars and P<0.01 and grows rate of Lv-shRNACdc42 cells compare to CON cells with three stars and P<0.001 was significant. after 96 and 120 h, this rate increased and grows rate of transduced cells compare to NC cells and CON cells with three stars and P<0.001 was significant.

## 4 Discussion

Cdc42 as downstream effector of EGFR, activated by phosphorylation and is involved in the regulation of critical cellular functions such as rearrangement of actin cytoskeleton, intracellular trafficking, cell polarity, cell-cycle regulation, cell fate determination and gene transcription. CDC42 is a key regulator involved in regulating the proliferation, migration and invasion of lung, breast, testicular, colorectal, esophageal and gastric cancer (Valdés-Mora et al. 2017; Du et al. 2015). Up-regulation of Cdc42 activates or inactivates some signaling pathways leading to create cancers. Cdc42 have been overexpressed in Non-Small Cell Lung Cancer (NSCLC) and progress cell cycle G1/S phases that make it a main target in cancer therapy (Li et al. 2013; Liu et al. 2011; Arias-Romero & Chernoff 2013; Qadir et al. 2015). In last 20 years, change in gene expression to target diseases, have been attracted many attention. Based on human genome sequencing and researches understands about molecular agents generating diseases, silencing pathogen genes is an attractive approach in treatment of wide range of diseases (Cejka et al. 2006). Gene therapy defined as the treatment of a disorder by the introduction of therapeutic genes into the appropriate cellular targets. These therapeutic genes can correct deleterious consequences of defected gene, or re-programme cell functions (Escors & Breckpot 2010). RNAi have many advantage compare to some treatment such as molecular drugs. RNAi technology have more specificity and wide range target capacity (Sakiragaoglu et al. 2013).

The determination of new cancer-specific targets and pathways amenable to RNAi therapy would provide a greater impetus to the promotion of targeted therapies. Several *in vitro* and *in vivo* proof-of-concept studies have mentioned that, RNAi has achieved from bench to bedside with several RNAi-based drugs in clinical trials (Bora et al. 2012). Inhibition of multiple genes is an effective approach to prevent or treat cancer as well as reduction of the possibility of multiple drugs’ resistance caused by overdose of chemical drugs (Mansoori et al. 2014). RNAi-based drugs with a gain of function genetic lesion or overexpression of disease causing gene(s), can be effective in the treatment of various diseases, such as cancer (Bora et al. 2012).

conducted on animal models emphasis that targeting essential proteins in the cell cycle, such as kinesin spindle protein (KSP) and polo-like kinase 1 (PLK1) by using specific siRNA have showed potent antitumor activity observed potent antitumor activity in both subcutaneous and hepatic tumor models (Mansoori et al. 2014). Judge *et al* showed that administration of siRNA targeting the essential cell-cycle proteins resulted in a 75% reduction in subcutaneous tumor size. Li *et al* demonstrated that an encapsulated siRNA specific to epidermal growth factor receptor (EGFR) were able to inhibit growth and metastasis in a mouse xenograft model of lung cancer.

Aleku *et al* showed that systemic administration a siRNA-lipoplex targeted in mice, rats and non-human primates, resulted in specific RNAi-mediated silencing of PKN3 expression and significant inhibition of tumor growth and lymph node metastasis.

Viral gene delivery systems consist of viruses that are modified to be replication-deficient which were made unable to replicate by redesigning which can deliver the genes to the cells to provide expression (Cevher et al. 2012).

Lentiviral vectors (LV) are efficient vehicles for gene transfer in mammalian cells due to their capacity to stably express a gene of interest in non-dividing and dividing cells and offer potential for treatment of a wide variety of syndromes including genetic/metabolic deficiencies, viral infection and certain types of cancers. Their use has exponentially grown in the last years both in research and in gene therapy techniques (Stengel & Zheng 2011).

In the current study, we transferred shRNA encoding lentiviruses into Calu6 cells as a cell line contributed to lung cancer. Generated recombinant lentiviruses analyzed for their cytotoxicity which results didn’t show any cytotoxicity of lentiviruses on the concentration which used for shRNA delivery.

Anti Cdc42 sequence carrying by lentiviruses had a reasonable potential to transfer shRNA and they could successfully target Cdc42 gene in Calu-6 cell line. shRNA mediated inhibition of Cdc42 gene led to reduction in the expression of Cdc42 gene which verified with real time PCR technique. Assessment the effect of Cdc42 gene expression reduction on proliferation in target cells showed significant decrease in calu-6 proliferation rate.

Our results were in accordance with other studies showing the effect of Cdc42 silencing on the inhibition of cell proliferation and cancer progression (Stengel & Zheng 2011).

In conclusion, our results proved that Cdc42 has critical role in proliferation of calu-6 cell lines and it may be a main target in lung cancer therapy but this subject should be considered that because of important roles of Cdc42 in other cellular processes, so silencing of this gene in *in vivo* experiments needed more control and specific targeted cancerous cells

## Abbreviation list

|  |  |
| --- | --- |
| **RNAi** | :RNA interference |
| **shRNA** | :short hairpin RNA |
| **Cdc42** | :Cell division control protein 42 |
| **GFP** | :green fluorescent protein |
| **MTT** | :3-(4,5-dimethylthiazol-2-yl)- 2,5-diphenyltetrazolium bromide |
| **MOI** | :Multiplicity Of Infection |
| **CT** | :Threshold Cycle |
| **CMV** | :Cytomegalovirus |
| **FACS** | :Fluorescence-Activated Cell Sorting |
| **DMSO** | :Dimethyl Sulfoxide |
| **OD** | :Optical Density |
| **LV** | :Lentiviral Vector |
| **CON** | :Control |
| **NC** | :Negative Control |

## Author contribution

Mohammad Nabiuni and Zohreh Ghambari: concept and design of the research; Zohreh Ghambari and Latifeh Karimzadeh: carrying out the experimental work and providing reagents; Zohreh Ghambari and Hanieh Jalali: data analysis and interpretation; Zohreh Ghambari and Latifeh Karimzadeh: writing of the article.

## References

Arias-Romero, L.E. & Chernoff, J., 2013. Targeting Cdc42 in cancer. Expert opinion on therapeutic targets, 17(11), pp.1263–73. Available at: http://www.ncbi.nlm.nih.gov/pubmed/23957315 [Accessed July 13, 2016].

Bolhassani, A. & Saleh, T., 2013. Challenges in Advancing the Field of Cancer Gene Therapy: An Overview of the Multi-Functional Nanocarriers. In Novel Gene Therapy Approaches. InTech.

Bora, R. et al., 2012. RNA interference therapeutics for cancer: Challenges and opportunities (Review). Molecular Medicine Reports, 6(1), pp.9–15. Available at: http://www.ncbi.nlm.nih.gov/pubmed/22576734 [Accessed May 29, 2017].

Cejka, D., Losert, D. & Wacheck, V., 2006. Short interfering RNA (siRNA): tool or therapeutic? Clinical Science, 110(1), pp.47–58. Available at: http://www.ncbi.nlm.nih.gov/pubmed/16336204 [Accessed January 21, 2017].

Cevher, E., Cevher, E. & Çağlar, E.Ş., 2012. Gene Delivery Systems: Recent Progress in Viral and Non-Viral Therapy. InTech.

Chou, M.M., Masuda-Robens, J.M. & Gupta, M.L., 2003. Cdc42 promotes G1 progression through p70 S6 kinase-mediated induction of cyclin E expression. The Journal of biological chemistry, 278(37), pp.35241–7. Available at: http://www.ncbi.nlm.nih.gov/pubmed/12842876 [Accessed March 14, 2015].

Dellaire, G., Berman, J.N. & Arceci, R.J., 2014. Cancer Genomics: From Bench to Personalized Medicine,

Donald D. Rao et al., 2009. siRNA vs. shRNA: Similarities and differences. Advanced Drug Delivery Reviews, pp.746–759.

Du, D. et al., 2015. Effects of CDC42 on the proliferation and invasion of gastric cancer cells. Molecular Medicine Reports, 13(1), pp.550–4. Available at: http://www.ncbi.nlm.nih.gov/pubmed/26549550 [Accessed May 29, 2017].

Escors, D. & Breckpot, K., 2010. Lentiviral vectors in gene therapy: their current status and future potential. Arch Immunol Ther Exp (Warsz), 58(2), pp.107–119.

Gómez Del Pulgar, T. et al., 2008. Cdc42 is highly expressed in colorectal adenocarcinoma and downregulates ID4 through an epigenetic mechanism. International journal of oncology, 33(1), pp.185–93. Available at: http://www.ncbi.nlm.nih.gov/pubmed/18575765 [Accessed September 7, 2015].

Humbert, O., Davis, L. & Maizels, N., 2012. Targeted Gene Therapies: Tools, Applications, Optimization. Crit Rev Biochem Mol Biol., 47(3), pp.264–281.

Li, Y. et al., 2013. miR-330 regulates the proliferation of colorectal cancer cells by targeting Cdc42. Biochemical and Biophysical Research Communications, 431, pp.560–565.

Liu, M. et al., 2011. miR-185 targets RhoA and Cdc42 expression and inhibits the proliferation potential of human colorectal cells. Cancer Letters, 301(2), pp.151–160. Available at: http://www.cancerletters.info/article/S0304383510005446/fulltext [Accessed September 7, 2015].

Mansoori, B., Sandoghchian Shotorbani, S. & Baradaran, B., 2014. RNA interference and its role in cancer therapy. Advanced pharmaceutical bulletin, 4(4), pp.313–21. Available at: http://www.ncbi.nlm.nih.gov/pubmed/25436185 [Accessed May 29, 2017].

Qadir, M.I., Parveen, A. & Ali, M., 2015. Cdc42: Role in Cancer Management. Chemical biology & drug design, 86(4), pp.432–9. Available at: http://www.ncbi.nlm.nih.gov/pubmed/25777055 [Accessed July 13, 2016].

Sakiragaoglu, O., Good, D. & Wei, M.Q., 2013. Cancer Gene Therapy with Small Oligonucleotides. In Novel Gene Therapy Approaches.

Song, H. & Yang, P.-C., 2010. Construction of shRNA lentiviral vector. North American journal of medical sciences, 2(12), pp.598–601. Available at: /pmc/articles/PMC3338230/?report=abstract [Accessed September 4, 2015].

Stengel, K. & Zheng, Y., 2011. Cdc42 in oncogenic transformation, invasion, and tumorigenesis. Cellular signalling, 23(9), pp.1415–23. Available at: http://www.pubmedcentral.nih.gov/articlerender.fcgi?artid=3115433&tool=pmcentrez&rendertype=abstract [Accessed March 14, 2015].

Valdés-Mora, F. et al., 2017. Clinical relevance of the transcriptional signature regulated by CDC42 in colorectal cancer. Oncotarget, 8(16), pp.26755–26770. Available at: http://www.ncbi.nlm.nih.gov/pubmed/28460460 [Accessed May 29, 2017].

